# Isolation of a Chlamydia Strain from Lung Tissue Samples in Healthy Chickens Exhibiting Antagonistic Effects on Avian Infectious Bronchitis Virus

**DOI:** 10.1101/2023.05.10.540153

**Authors:** Miaoxiao Zhen

## Abstract

To assess the feasibility of isolating Chlamydia strains with antiviral activity against avian infectious bronchitis virus (IBV) from healthy chickens in farms, 58 Chlamydia strains were obtained from 160 lung tissue samples collected across ten chicken farms, yielding an overall isolation rate of 36.25%. SPF chickens co-infected with Chlamydia and IBV were employed for verification, leading to the identification of eight strains (B_i_ ≥ 0.800) with potent antiviral effects against IBV, accounting for 13.79% of the total isolated strains. The top-performing strain, Y17 Chlamydia strain, was selected and subjected to cell co-culture and U-tube experiments with IBV. Results demonstrated that the Y17 Chlamydia strain significantly impeded IBV replication in chicken tracheal epithelial cells (P<0.01) and did not secrete or induce host cells to secrete extracellular metabolites with antagonistic effects on IBV infection in U-tube experiments (P>0.05), suggesting that its antiviral activity was cell-autonomous. Our research highlights the feasibility of isolating Chlamydia strains with antiviral activity against IBV from healthy chickens and suggests that antiviral strains could be widespread, even though their efficacy against viruses is strain-specific. The presence of broad-spectrum antiviral Chlamydia strains may also be possible. Considering the prevalence of Chlamydia strains in vertebrate hosts, along with the fact that some strains are either non-pathogenic or display low toxicity, our findings could offer a fresh perspective on the prevention and treatment of viral infections in vertebrates.

## 0 Introduction

Interspecies competition among various bacteria can lead to secondary metabolite production, from which many antibiotics are derived and extracted^[1,2,3]^. Notably, the majority of natural antibiotics are obtained from actinomycetes’ secondary metabolites, particularly the Streptomyces genus^[4,5]^. These microorganisms share features of both bacteria and fungi. Chlamydia are intracellular parasites that exist between bacteria and viruses, relying on their host cells for essential high-energy compounds like adenosine triphosphate (ATP) and adenosine diphosphate (ADP)^[6,7]^. These organisms possess a biphasic life cycle, including infectious elementary bodies (EB) and non-infectious reticulate bodies (RB)^[6,7]^. Chlamydia relies on a type III secretion system to release products that regulate host cell metabolism, providing necessary resources and energy for their growth and proliferation^[8,9]^. Viruses, similar to Chlamydia, are another small obligatory intracellular parasites that depend on host cell resources for replicative needs^[10]^. Given Chlamydia and viruses reside within the same host cell niches, they may compete directly, raising the possibility of isolating virus-resistant Chlamydia strains^[11]^.

Various studies have indicated that certain Chlamydia strains may provide antiviral protection in hosts^[12-24]^. For instance, Hongtao et al. utilized electron microscopy to detect large quantities of Chlamydia-like factors in organs, such as the lungs, spleen, liver, kidney, and lymph nodes of some severe acute respiratory syndrome (SARS) patients, with few to no SARS-CoV particles observed^[12]^. Further results indicated that only a small percentage of Chlamydia-like factors isolated by Hongtao et al. reacted with antibodies present in SARS patients’ blood (less than 10%), suggesting they were not SARS pathogens but instead normal microorganisms within their host^[13]^. Slade et al. reported that *Chlamydia muridarum* (Cm) Wiess strain could protect mice from herpes simplex virus 2 (HSV-2) induced neural damage, and this protective effect was unrelated to host interferon-beta (IFN-*β*) induction by Chlamydia inoculation^[14,15]^. Alosaimi et al. discovered that co-infection with influenza virus and SARS-CoV-2 increased the likelihood of intensive care unit (ICU) monitoring for COVID-19 patients, while co-infection with bacteria, including *Chlamydia pneumoniae* (*C. pneumoniae*), lessened this probability^[16]^. Mina et al. reanalyzed PCR detection data from respiratory disease patients in Oslo, Norway and found a strong antagonistic relationship between *C. pneumoniae* infection and influenza virus infection^[17]^. Similar results were observed in two other sufficiently sampled studies conducted by Reinton et al. and Lieberman et al.^[18,19]^. Prusty et al. identified a significant negative correlation between *C. trachomatis* load and Human Herpesvirus-6 (HHV-6) viral load in co-infected patients^[20]^.

Borel et al. found that *Chlamydia pecorum* 1710S strain, isolated from pigs, could significantly inhibit syncytial formation of Porcine Epidemic Diarrhea Virus (PEDV) infected Vero-6 cells; however, the *Chlamydia abortus* S26/3 strain, isolated from sheep, did not exhibit the same reduction^[21]^. Superti et al. observed that co-infection of *Chlamydia trachomatis*(*C. trachomat*)serovar E BOUR strain and HSV-2 333 strain significantly decreased HSV-2 333 strain replication in HT-1376 cells, while the co-infection of *C. trachomatis* increased HSV-2 replication in HeLa-229 cells^[22,23]^. Erin et al. identified a significant negative correlation between *Chlamydia pecorum* infection and Burpengary virus infection in South Australian koalas^[24]^. Wolbachia, a widespread parasitic or symbiotic bacterium found in arthropods and some nematodes, exhibits characteristics intermediate between bacteria and viruses^[25]^. Some Wolbachia strains could shorten the life of *Drosophila melanogaster*^[26,27]^Recent studies have revealed that some symbiotic strains can substantially reduce the viral load of multiple RNA viruses in their hosts, including Dengue virus, Norovirus, and Yellow Fever virus, and even extend the host’s lifespan. Different strains display antagonism against different viruses in the host, and some strains can even combat multiple viruses^[25,26,28]^.

Though some studies propose that co-infection of specific Chlamydia strains with relevant pathogenic viruses may worsen host conditions and reduce survival rates, this might be attributed to the Chlamydia strain, virus strain, infection sequence, and infection time interval ^[14,15,21,25,27,30,31,32,33,34,35,36,37]^. Various Chlamydia strains and virus strains may display different interactions in the host. Evidently, whether synergistic or unrelated, the co-infection of Chlamydia strains and relevant virus strains offers no benefit to the host, and only antagonistic relationships may assist the host in resisting viral infections. Chlamydia is widely distributed in nature, and individual strains exhibit varying pathogenicity. Humans carry various Chlamydia strains, with only a small number being pathogenic^[38,39,40]^. Heike et al. reported that the Chlamydia psittaci carriage rate in tits (Paridae) in Germany was 54.3%, and only strains present in diseased tits showed cytopathic effects (CPE) when inoculated into BGM cells^[41]^. Mohamad et al. demonstrated an evolutionary trend from high to low virulence in Chlamydia pecorum strains in Australian koalas ^[42]^. Thus, in addition to focusing on pathogenic Chlamydia strains, attention should be given to non-pathogenic Chlamydia strains that parasitize the host and may provide benefits, such as aiding in host defense against pathogenic viruses or bacteria^[14,22,25,43,44]^.

Considering the above factors, the following experiments aim to isolate non-pathogenic or low virulence Chlamydia strains, opposing Infectious Bronchitis Virus (IBV) from lung tissue samples obtained from ten chicken farms in Guangzhou, China. The experiments will preliminarily determine whether the isolated Chlamydia strains secrete or induce host cells to secrete metabolites antagonizing IBV. The choice of isolating Chlamydia strains from healthy chickens and using IBV as the validation virus strain is based on the following rationale: (i) the wide availability of materials for isolating Chlamydia strain. Some previous studies have shown that there is widespread Chlamydia infection in chicken farms across China^[45,46]^ ; (ii) IBV is harmless to humans, and evidence suggests that IBV infection is widespread in various regions of China^[47,48]^. Hence, it is more practical to assess the antagonistic ability of isolated Chlamydia strains. (iii)Specific pathogen-free (SPF) chicks, which are widely available and cost-effective, serve as suitable subjects for verification experiments. (iv) IBV serves as a prototype virus for the γ-coronavirus genus within the Coronaviridae family. Success in this area may offer valuable insights for developing novel methods to combat SARS-CoV-2^[49]^.

## 1 Materials and Methods

### 1.1 Materials, Reagents, and Instruments

Our study involved lung tissue samples from 160 adult healthy chickens from ten chicken farms in Guangzhou, China; IBV strain (Number: CVCC AV1511) and avian influenza virus H_9_N_2_ strain (AIV, Number: CVCC AV1554) purchased from the China Veterinary Microbes Culture Collection Center; 1500 specific pathogen-free (SPF) chicks (14-day-old, from Xin Xing Da Hua Poultry Egg Co., Ltd.); chicken tracheal epithelial cells (obtained from Wuhan Puno Life Science Technologies Co., Ltd.); and BGM cells (sourced from the Laboratory of Animal Biotechnology and Immunology, Ghent University).

Key reagents included fetal bovine serum (FBS) and Dulbecco’s Modified Eagle Medium (DMEM) culture medium (GIBCO); Viral RNA kit (QIAGEN, Germany); Reverse Transcription Kit (Promega, USA); IBV nucleic acid amplification fluorescence detection kit (Shanghai Furex Medical Scientific and Technological Development Co., Ltd.); CellTox Green Cytotoxicity Assay Kit (Promega, USA); and SYBR Green PCR Master Mix Kit (Baosheng Bioscience (Dalian) Co., Ltd.)

The main instruments used were a cell culture incubator; PE5700 automatic fluorescent quantitative PCR analyzer (ABI, USA); CKX41 inverted microscope (Olympus, Japan); U-tube; 0.2um microfiltration membrane, 0.05um ultrafiltration membrane, 0.002um nanofiltration membrane (Xiamen Fumei Technology Co., Ltd.); and 13 mm round cover glass.

### 1.2 Isolation of Chlamydia Strains

Chlamydia strains were isolated from lung tissue samples obtained from 160 adult healthy chickens from ten chicken farms in Guangzhou, China. After a series of sample preparations, inoculations, and detections, the Chlamydia inclusion bodies were observed under a microscope. Chlamydia strains were stored in sucrose-phosphate-glutamate (SPG) solution at -80°C for future use. The specific steps are as follows:

i. Collect lung tissue samples from 160 adult healthy meat chickens originating from 10 chicken farms in Guangzhou, China. Quickly place the samples in sampling tubes and transport them to the laboratory at low temperatures. Add a 4-fold volume of sterilized PBS solution and use a glass homogenizer to grind the tissues until no obvious chunks remain. Centrifuge the samples at 3000 r/min at 4°C for 5 minutes and collect the supernatant. Add 100 μg/mL gentamicin and 100 μg/mL streptomycin sulfate for bacteria removal through incubation at room temperature for 1 hour, followed by centrifugation for 20 minutes at 4°C, 3000 r/min. Store the samples at -80°C for later use.
ii. Inoculate BGM cells in culture tubes at a density of 1 × 10^⋀4^cells/tube using 8% PBS DMEM culture medium, which additionally includes 100 μg/mL fetal bovine serum, 292 μg/mL glutamine, 10 g/L vitamin solution, 100 μg/mL gentamicin sulfate solution, 100 μg/mL streptomycin sulfate solution, and 2 μg/mL fungizone. Place 13 mm round cover glasses in the tubes and, once the cell density reaches 80%, add 100 μL of sample per tube. Incubate the tubes in a cell culture box at 37°C, 5% CO_2_ concentration, and full humidity for 72 hours. Perform three replicates per sample and blindly passage three generations, while also setting a negative control.
iii. Remove the 13 mm round cover glass, fix it with ethanol, and perform Giemsa staining. Observe Chlamydia inclusion bodies under a microscope at 400x magnification^[50]^. Once positive results are confirmed, add 1 mL of sucrose-phosphate-glutamate (SPG) solution (218 mmol/L sucrose, 38 mmol/L KH_2_PO_4_, 7 mmol/L K_2_HPO_4_, and 5 mmol/L glutamic acid) to the remaining culture tubes and store them at -80°C for future use.

### 1.3 SPF Chicks (14-day-old) Co-infection Experiment with Chlamydia and IBV

We conducted a co-infection experiment using SPF chicks and the isolated Chlamydia strains by inoculating them in different sequences. Throat swab samples were collected on the 6th, 9th, 12th, and 15th days after the first inoculation to measure Chlamydia and IBV levels in the chicks. The mortality rate for each group, consisting of 10 chickens per group, was calculated. The specific steps are as follows:

i. Prepare an inoculation solution for each isolated Chlamydia strain from chickens at a concentration of 1 × 10^7^ IFUs/mL, and inoculate it into the trachea of SPF chicks.
ii. Prepare an inoculation solution containing the IBV strain and dilute it to a concentration of 5 × 10^4^ PFU/mL. Inoculate the SPF chicks three days after the Chlamydia inoculation.
iii. On the 6th, 9th, 12th, and 15th days following the initial inoculation, detect shedding levels of Chlamydia and IBV in the throat swab samples of the chicks, and calculate the mortality rate for each group of 10 SPF chicks.
iv. Establish a negative control group with Chlamydia and IBV inoculations. As depicted in Figure 1(a).

**Figure 1.**
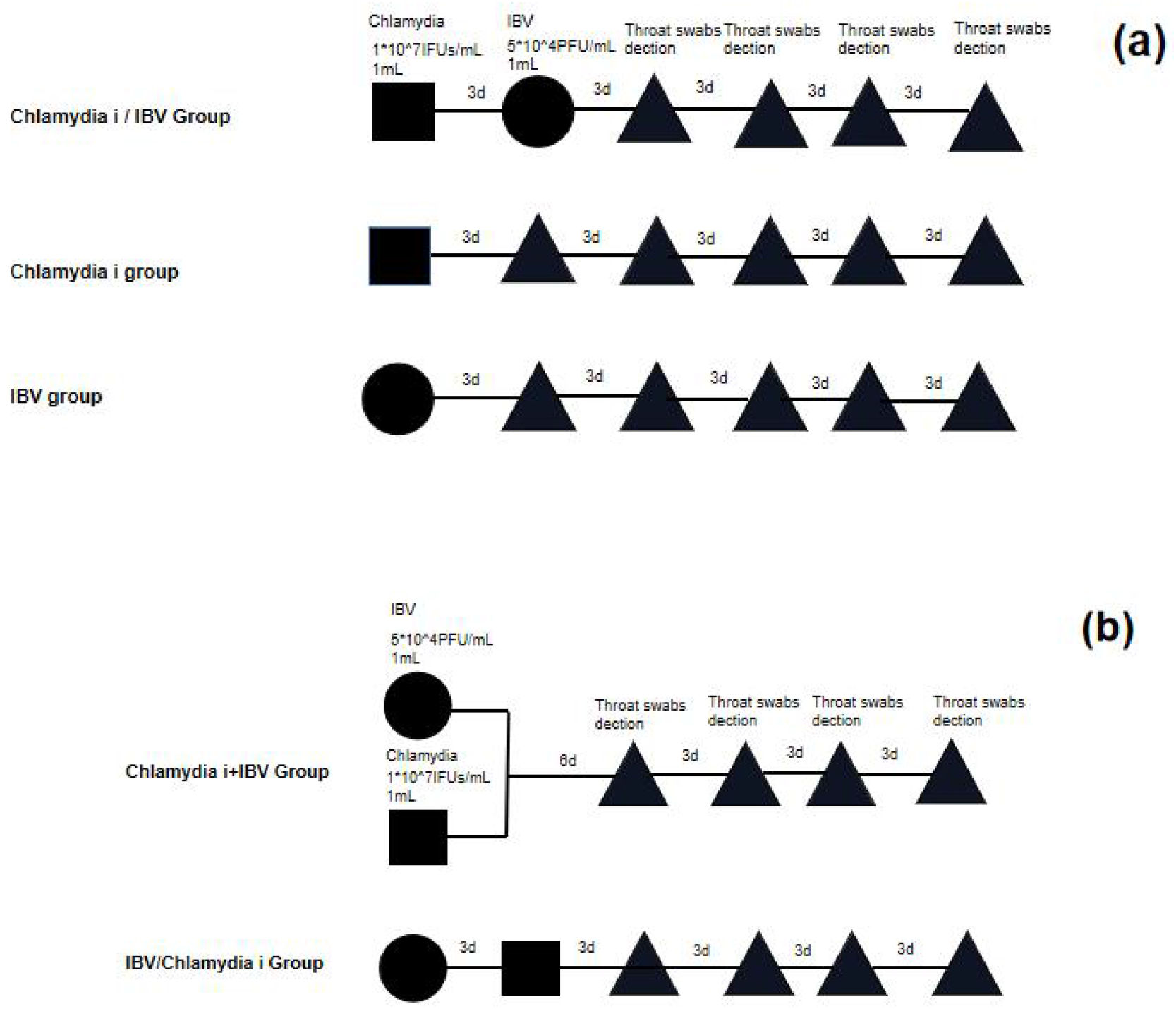
illustrates the inoculation sequence and detection time for the co-infection of Chlamydia and IBV in SPF chicks. Figure 1(a) presents the inoculation sequence and detection time for the group receiving Chlamydia first, followed by IBV (Chlamydia i/IBV group) and control groups (Chlamydia i group and IBV group). Figure 1(b) depicts the inoculation sequence and detection time for the group simultaneously receiving Chlamydia and IBV (Chlamydia+IBV group) and the group receiving IBV first, followed by Chlamydia (IBV/Chlamydia group).

The calculation score for the i-th isolated Chlamydia strain is obtained by calculating the mortality rate of SPF chicks in the Chlamydia group, the mortality rate of SPF chicks in the IBV inoculation group, the mortality rate of SPF chicks in the mixed inoculation group, and the shedding levels of Chlamydia and IBV in the throat swab samples of the mixed inoculation group, according to equation 3-1 and then equation 3-2.

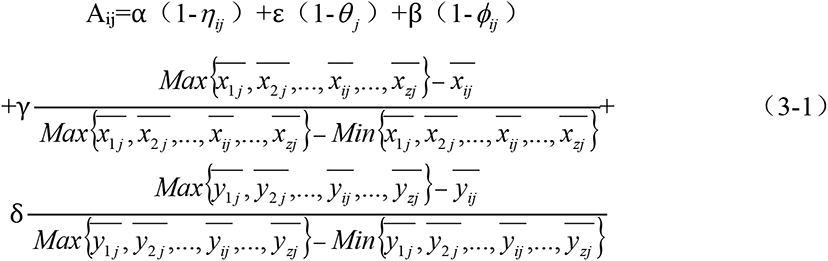

In the formula, i denotes the i-th strain of isolated Chlamydia; Z signifies the total number of isolated Chlamydia strains; j represents the j-th detection; A_ij_ corresponds to the score of the i-th bacterial strain after the j-th detection;*η*_*ij*_ standsfor the SPF chicken mortality rate in the Chlamydia inoculation group for the i-th strain during the j-th detection;*θ*_*j*_ indicates the SPF chicken mortality rate in the IBV inoculation group during the j-th detection; *ϕ*_*ij*_ signifies the mortality rate of SPF chickens in the mixed inoculation group of the i-th strain during the j-th detection;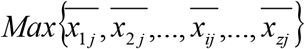represents the maximum Chlamydia shedding levels in each mixed inoculation group at the j-th detection; 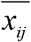 pertains to the Chlamydia shedding levels in the mixed inoculation group of the i-th bacterial strain during the j-th detection of throat swab specimens;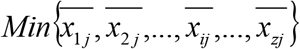 denotes the minimum Chlamydia shedding levels in each mixed inoculation group at the j-th detection; 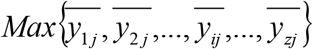 corresponds to the maximum IBV shedding levels in each mixed inoculation group during the j-th detection; 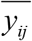 illustrates IBV shedding levels in the throat swab samples of the mixed inoculation group of the i-th Chlamydia strain during the j-th detection;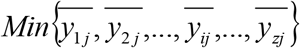 embodies the minimum IBV shedding levels in each mixed inoculation group at the j-th detection; α, β, γ, δ, ε respectively represent assignment coefficients, and the sum of coefficients equals 1; in this study, we will adopt α and ε coefficient values of 0.2, β coefficient values of 0.3, and γ and δ coefficient values of 0.15.

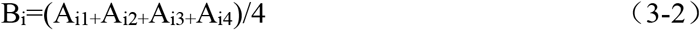

In the formula, B_i_ denotes the calculated score for the i-th Chlamydia strain, while A_i1_, A_i2_, A_i3_, and A_i4_ signify the individual scores from each detection (on the 6th, 9th, 12th, and 15th days) for the i-th Chlamydia strain. The top three Chlamydia strains, as determined by their highest B_i_ scores, were selected for further investigation. These strains were inoculated into SPF chicks using two methods: simultaneous inoculation and a sequential approach, consisting of IBV inoculation first, followed by Chlamydia inoculation three days later. Throat swab samples were collected from the SPF chickens on the 6th, 9th, 12th, and 15th days after initial inoculation to assess shedding levels of Chlamydia and IBV. Each group’s mortality rate, comprising 10 chickens per group, will be determined, as illustrated in Figure 1(b). The final scores for the three Chlamydia strains will be calculated according to formula 3-3, and the strain with the highest C_i_ score will be chosen for additional experimentation.

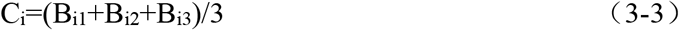

In the formula, C_i_ denotes the final score for the i-th Chlamydia strain; B_i1_, B_i2_, and B_i3_ signify the scores for the i-th Chlamydia strain when inoculated via three distinct methods -concurrent inoculation, initial Chlamydia inoculation followed by IBV inoculation, and primary IBV inoculation succeeded by Chlamydia inoculation -in SPF chickens as computed using formulas 3-1 and 3-2.

### 1.4 Cell Co-cultivation and U-shaped Tube Experiment of Chlamydia and IBV

#### 1.4.1 Cell Co-cultivation of Chlamydia and IBV

Previous studies have demonstrated that interactions between Chlamydia and viruses in cultured cells may vary based on the specific cell lines, producing opposing results in different culture cell lines^[22,23]^. To more accurately represent the real situation, we utilized chicken tracheal epithelial cells for co-culturing Chlamydia strains and IBV. The Chlamydia strain with the highest C_i_ score and IBV were inoculated in three ways: simultaneous inoculation, inoculation of Chlamydia followed by IBV, and inoculation of IBV followed by Chlamydia, onto chicken tracheal epithelial cells. A 13mm glass coverslip was placed, and the cell inoculum was 1^*^10^4^ cells/flask (the cell culture medium was obtained from Wuhan PunoLife Biotechnology Co., Ltd., and the culture method was performed according to their instructions). The cells were generally inoculated with infectious liquid when the cell density reached 80%. The inoculation time interval was 24 hours. The levels of Chlamydia and IBV in each group were measured at 72 hours after the first inoculation. A negative control group (inoculation groups for Chlamydia strain and IBV strain) was also established. Each group was replicated three times. The specific experimental process is depicted in Figure 2(a).

**Figure 2.**
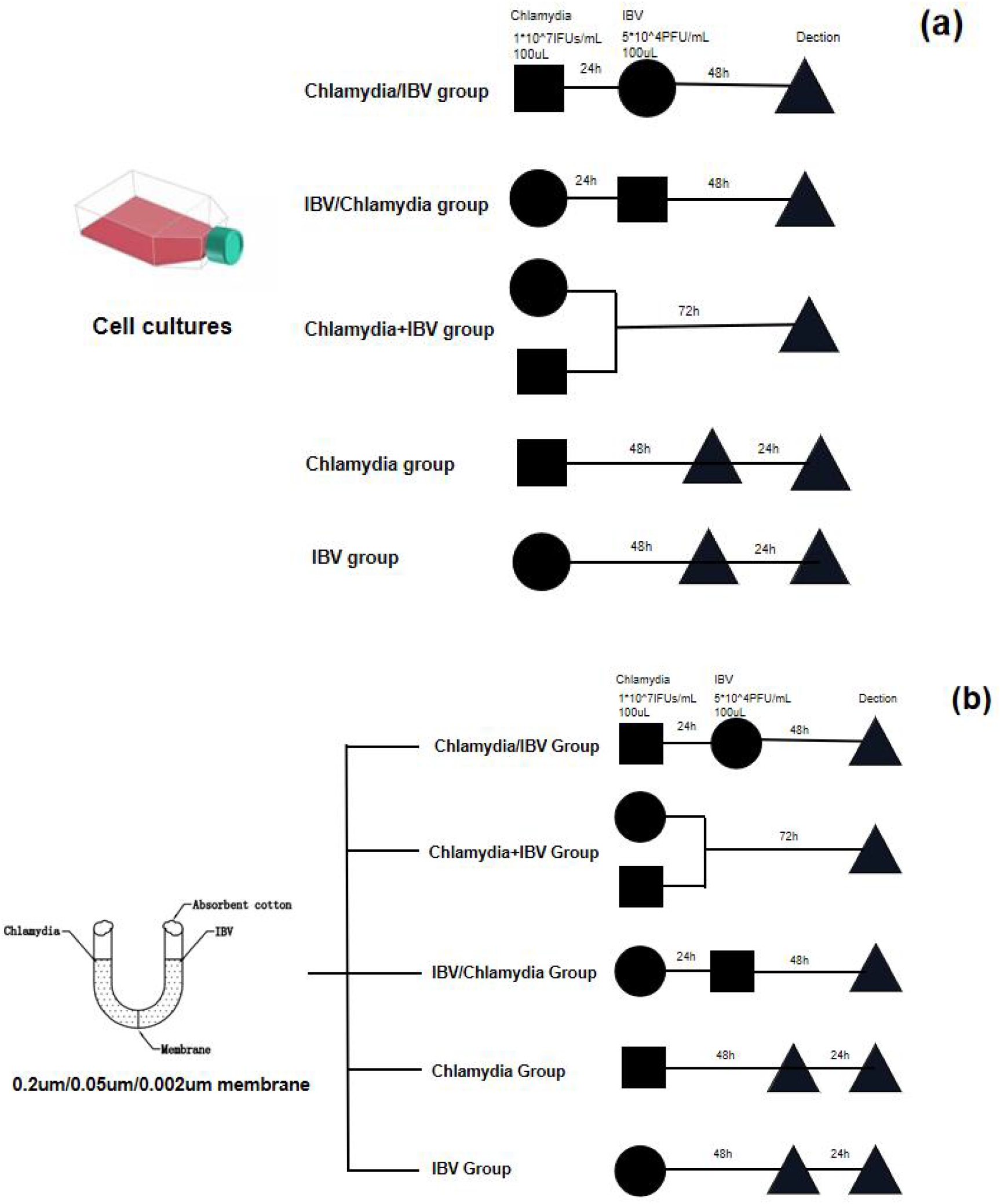
displays the cell co-cultivation with Chlamydia and IBV in chicken tracheal epithelial cells, as well as the inoculation sequence of Chlamydia and IBV inoculation in U-tube experiments. Figure 2a represents the inoculation sequence and detection time for the groups inoculated with Chlamydia first and then IBV (Chlamydia/IBV), simultaneously with Chlamydia and IBV (Chlamydia+IBV), IBV first and then Chlamydia (IBV/Chlamydia), and the control groups (Chlamydia group and IBV group) in cell co-cultivation. Figure 2b exhibits the inoculation sequence and detection time for the Chlamydia/IBV, Chlamydia+IBV, IBV/Chlamydia, and control groups (Chlamydia group and IBV group) in U-tube experiments with three pore sizes membrane (0.002um/0.05um/0.2um).

#### 1.4.2 U-shaped Tube Experiment

A 0.2μm microfiltration membrane, a 0.05μm ultrafiltration membrane, and a 0.002μm nanofiltration membrane were positioned in the center of the U-shaped tube. For each pore size, Chlamydia (on the left) and IBV (on the right) were inoculated simultaneously; Chlamydia was first introduced on the left, followed by IBV inoculation on the right; IBV was initially inoculated on the right, followed by Chlamydia on the left. A 13mm glass cover slip was placed on both sides of the U-shaped tube, with equal volumes of cell culture medium on each side, and a cell inoculum of 1^*^10^4^ cells. As the cell density increased to 80%, the infection liquid could be introduced. The inoculation time interval was set at 24 hours, and Chlamydia and IBV levels in each group were measured 72 hours after the initial inoculation. Each test group was performed in triplicate, and a negative control group was also included. The detailed experimental process is illustrated in Figure 2(b).

### 1.5 Detection of Chlamydia

A 13mm cover slip was removed and stained with Giemsa. The number of inclusions within the field of view was counted using a 400x magnification microscope, and the Chlamydia level was determined utilizing formulas 3-4.

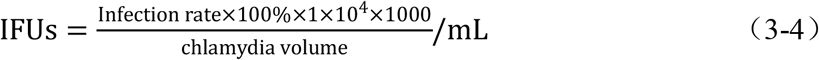

In the formula, IFUs represent the number of active Elementary Bodies (EBs) capable of forming inclusion bodies per milliliter. The infection rate refers to the ratio of the number of inclusion bodies observed under a microscope to the total number of cells. 1*10^4^ denotes the total number of cells, while 1000 serves as the conversion unit (uL-mL); the chlamydia volume signifies the volume of added chlamydia infective fluid.

### 1.6 Detection of IBV

Viral RNA was extracted using a Viral RNA kit, and reverse transcription was performed with a Reverse Transcription Kit to obtain IBV cDNA. An automated fluorescent quantitative PCR analyzer was employed to amplify the cDNA and detect the IBV levels. The 20 uL qRT-PCR reaction mixture included 2 uL cDNA, 10 uL 2×SYBR Green qPCR Mix, 0.4 uL forward primer, 0.4 uL reverse primer, and 7.2 uL RNAase-free water. Reaction conditions were set at 55°C for 10 min, 95°C for 3 min, followed by 45 cycles of amplification at 95°C for 10 s, and then held at 60°C for 34 s. IBV primers were referenced from Calisson et al.^[51]^and synthesized by Sangon Biotech (Shanghai) Co., Ltd., with the following primer sequences: F: 5’-GCTTTTGAGCCTAGCGTT-3’; R: 5’-CCATGTTGTCACTGTCTATTG-3’. AIV H_9_N_2_ strain primer sequences were taken from Chu Jun^[31]^ and synthesized by Sangon Biotech (Shanghai) Co., Ltd., with the primer sequences as follows: F:5’-CTGGAATCTGAGGGAACTTACAAAA-3’; R:5’-GAAGGCACAAACCCCATT-3’.

### 1.7 Statistical Analysis

After calculating the mean values of the detection for each group of Chlamydia strains and IBV co-cultured with SPF chicks, a comparison analysis was conducted. The detection data from cell co-cultivation and U-tube experiments were presented in the form of mean ± standard deviation (SD) and compared using t-student two-tailed tests and one-factor variance analysis (ANOVA). The analysis software used was SPSS 25.0. Generally, results are considered not significant when p > 0.05; significant when p < 0.05; and extremely significant when p < 0.01.

## 2 Results

### 2.1 Isolation Results of Chlamydia Strains

We isolated 58 Chlamydia strains from 160 healthy chicken lung tissue samples, achieving an overall isolation rate of 36.25%, as shown in Table 1.

**Table 1.**
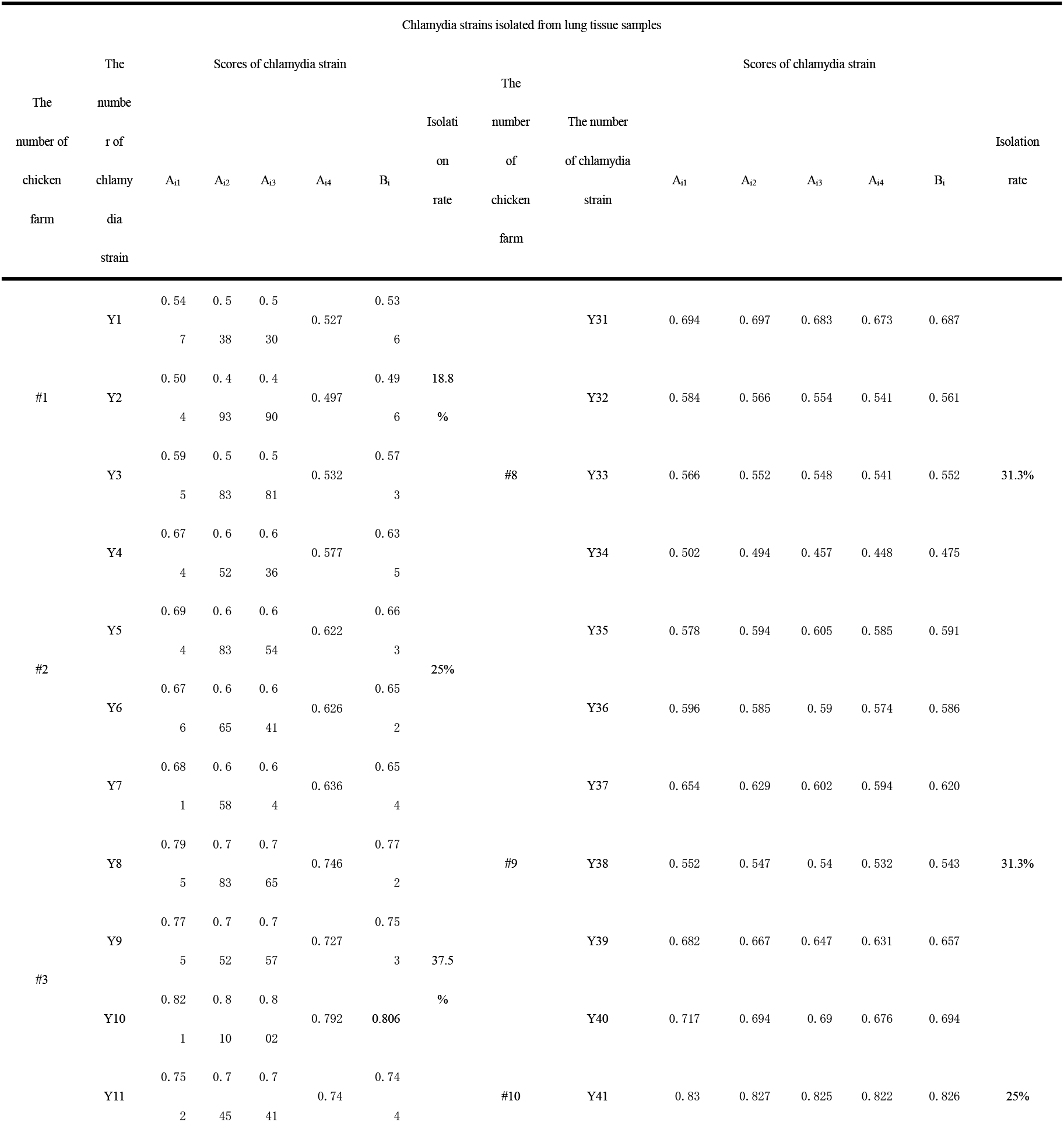
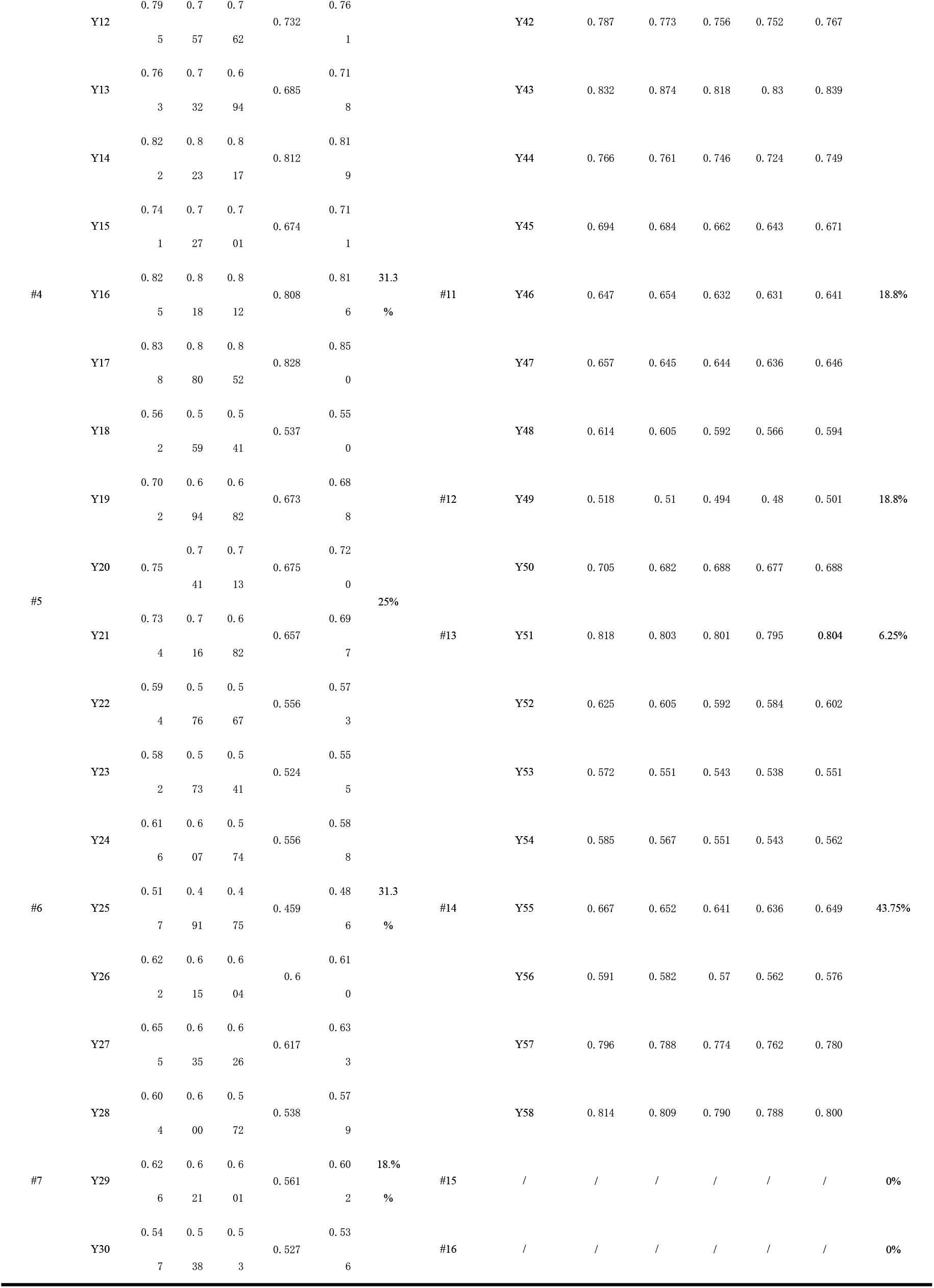
Statistics on the number and score (A_i_ and B_i_) of Chlamydia strains isolated from healthy chickens in ten chicken farms.

### 2.2 Infection Experiment Results of Chlamydia and IBV in SPF Chickens (14 days old)

As per Table 1, the overall scores of these isolated Chlamydia strains ranged from 0.716 to 1.030. With B_i_ ≥ 0.800 as the boundary, we considered the isolated Chlamydia strains to exhibit a good inhibitory effect on IBV. We isolated eight Chlamydia strains with a strong inhibitory effect on IBV, accounting for 5% of the total isolated samples and 13.79% of the Chlamydia isolates. The three strains with the highest scores were Chlamydia strains Y17, Y41, and Y43, with Y41 and Y43 originating from the same chicken farm. Table 2 shows that the Y17 Chlamydia strain had the highest C_i_ score. Hence, we utilized it for subsequent cell co-cultivation and U-tube experiments. As demonstrated in Figure 3, there was no significant difference in Chlamydia shedding levels between the Chlamydia/IBV group, Chlamydia+IBV group, IBV/Chlamydia group, and Chlamydia group in the four Chlamydia shedding levels detection of the Y17 Chlamydia strain (P>0.05). The IBV shedding levels of the Chlamydia/IBV group, Chlamydia+IBV group, and IBV/Chlamydia group were significantly lower compared to the IBV group (P<0.01). By the 15th day, SPF chickens’ survival rates in the Chlamydia/IBV group, Chlamydia+IBV group, IBV/Chlamydia group, Chlamydia group, and IBV group were 100%, 100%, 90%, 100%, and 60%, respectively. Consequently, there was no significant difference in SPF chickens’ survival rates between the Chlamydia/IBV group, Chlamydia+IBV group, IBV/Chlamydia group, and Chlamydia group (P>0.05). However, a significant difference existed between these groups and the IBV group (P<0.01). These data suggest that the Y17 Chlamydia strain provides significant protection against IBV infection in SPF chickens and might exhibit low virulence to the host SPF chickens.

**Table 2.**
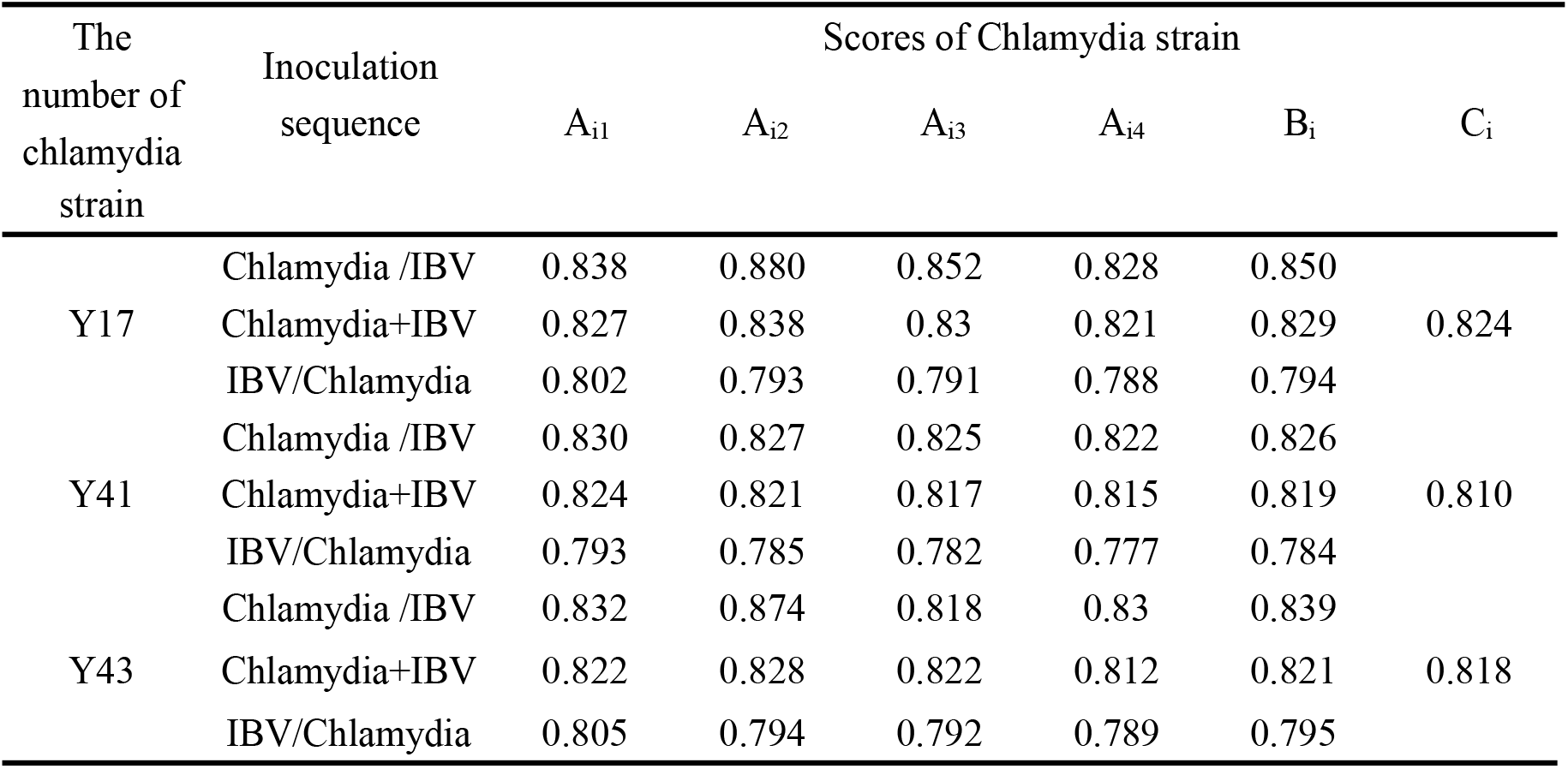
Scores (C_i_) obtained from SPF chicks infected with three Chlamydia strains.

**Figure 3.**
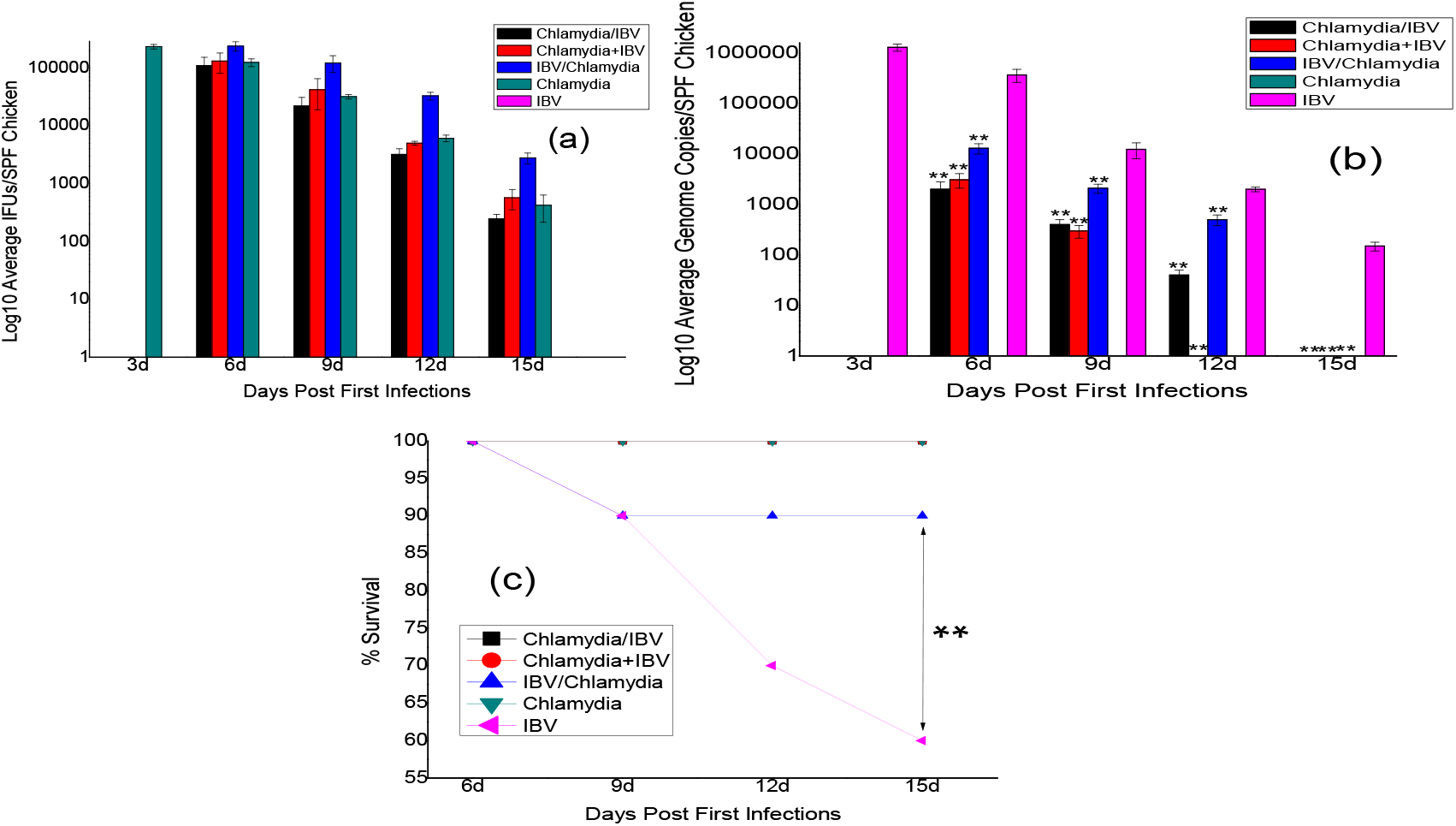
demonstrates the outcomes of co-infection between the Y17 Chlamydia strain and IBV in SPF 14-day-old chicks. Figure 3(a) shows the changes in the shedding levels of Y17 Chlamydia strain for the Chlamydia/IBV, Chlamydia+IBV, IBV/Chlamydia, Chlamydia, and IBV groups. Figure 3(b) displays the changes in the shedding levels of IBV for the same groups. Figure 3(c) exhibits the survival rate of the SPF chicks in these groups.

### 2.3 Cell Co-culture Results of Chlamydia and IBV

Following 72 hours of culture, we observed normal cell growth in the Chlamydia/IBV group, Chlamydia+IBV group, IBV/Chlamydia group, and Chlamydia group, with no evident CPE. In the control group (IBV group), CPE occurred in the cells, suggesting that the isolated strains might be harmless to the host. This finding is consistent with Heike et al.’s results, which indicated when Chlamydia strains isolated from healthy wild tits were inoculated into BGM cells, the BGM cells did not exhibit CPE^[41]^. As shown in Figure 4, our cell co-culture results revealed no significant differences in Chlamydia levels between the Chlamydia/IBV group, Chlamydia+IBV group, and Chlamydia group (P>0.05). However, the Chlamydia levels of the IBV/Chlamydia group were significantly lower than the Chlamydia group (P<0.01). This finding may be related to IBV pre-infection, which damaged the host cell surface receptor or altered the host cell metabolic pathway, leading to decreased proliferation of Chlamydia strains^[22]^. The IBV levels for the Chlamydia/IBV group, Chlamydia+IBV group, and IBV/Chlamydia group were considerably lower than the IBV group (P<0.01), indicating that the Y17 Chlamydia strain significantly reduced IBV replication in chicken tracheal epithelial cells. We also cultured chicken epithelial cells with the Y17 Chlamydia strain and AIV, and similar results was obtained, implying that the antiviral effect of this Chlamydia strain might be broad-spectrum (Figure 1S).

**Figure 4.**
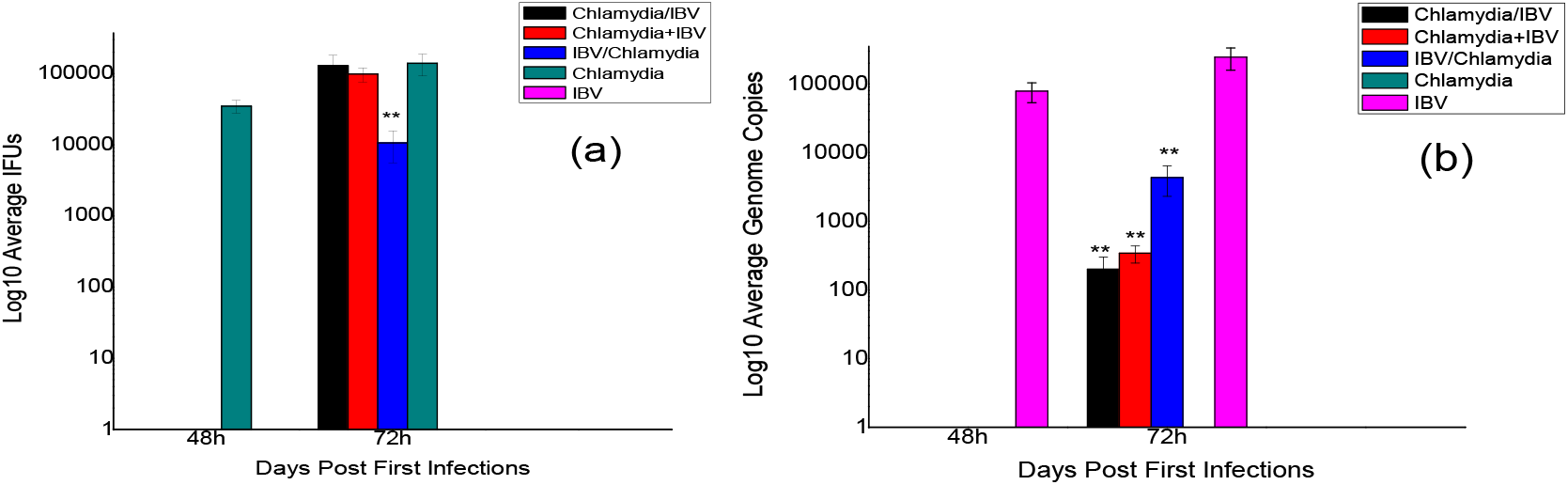
presents the results of cell co-cultivation with Chlamydia and IBV in chicken tracheal epithelial cells. Figure 4(a) highlights the changes in Chlamydia levels of groups, including Chlamydia/IBV, Chlamydia+IBV, IBV/Chlamydia, Chlamydia, and IBV. Figure 4(b) depicts the changes in IBV levels of these same groups.

### 2.4 U-tube Experiment Results

Figure 5 demonstrates no significant difference in Chlamydia levels between the Chlamydia/IBV group, Chlamydia+IBV group, IBV/Chlamydia group, and Chlamydia group under the 0.2um, 0.05um, and 0.002um pore size membranes (P>0.05). This observation indicates that the proliferation of IBV does not impact the cellular proliferation of the Y17 Chlamydia strain. There were no significant differences in IBV detection values between the Chlamydia/IBV group, Chlamydia+IBV group, IBV/Chlamydia group, and IBV group under the 0.2um, 0.05um, and 0.002um pore size membranes (P>0.05). This outcome suggests that the Y17 Chlamydia strain does not induce the secretion of extracellular metabolites that inhibit IBV replication, and its antiviral effect is cell-autonomous, consistent with Nainu et al.’s findings on Wolbachia antiviral protection in insects^[52]^.

**Figure 5.**
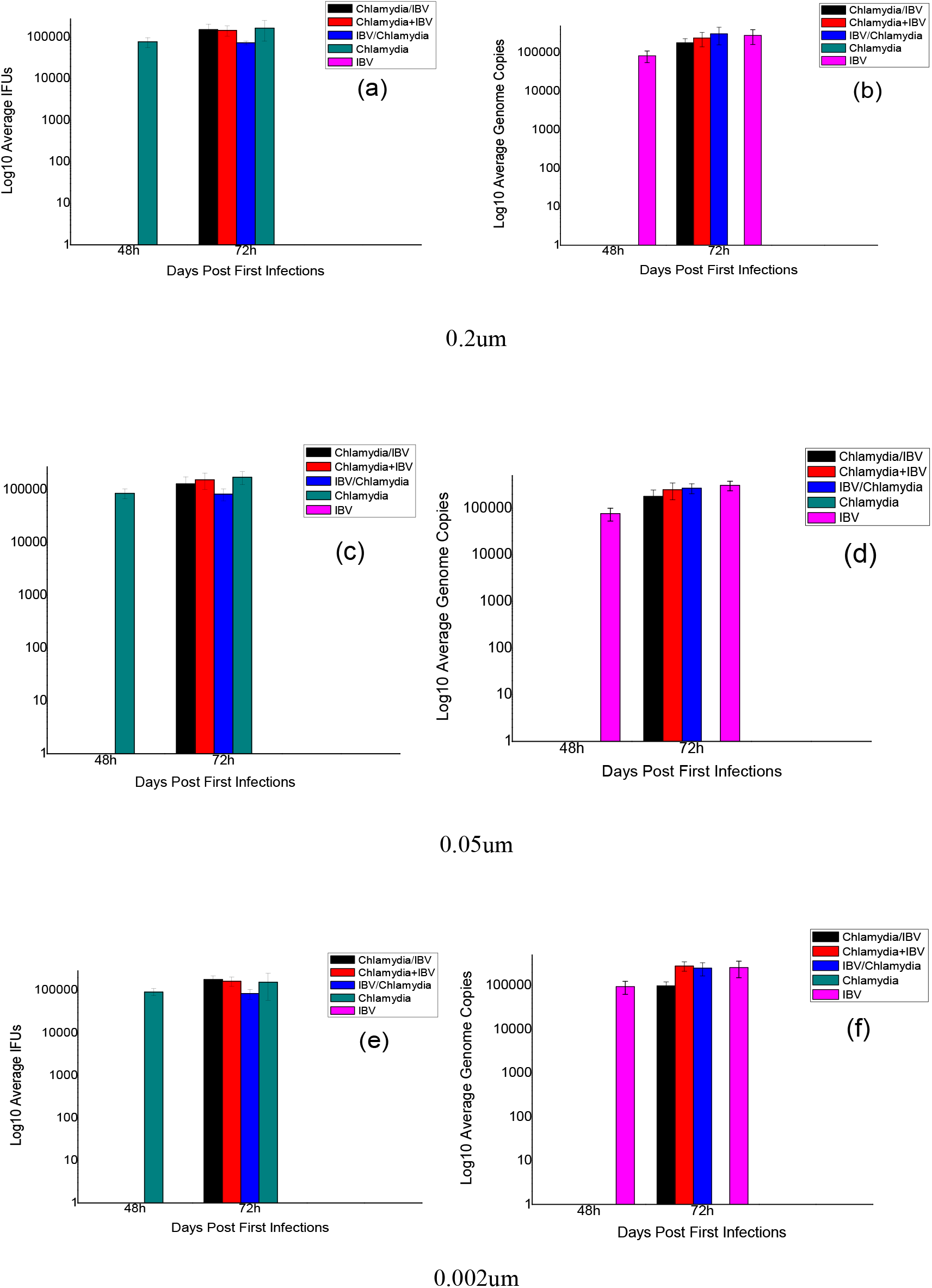
reveals the results of the U-tube experiment with Chlamydia and IBV. Figure 5(a) displays the change in Y17 Chlamydia strain levels for the Chlamydia/IBV group, Chlamydia + IBV group, IBV/Chlamydia group, Chlamydia group, and IBV group under the 0.2um pore size membrane; Figure 5(b) shows the change in IBV levels for the same groups under the 0.2um pore size membrane. Figure 5(c) presents the change in Y17 Chlamydia strain levels for the same groups under the 0.05um pore size membrane; Figure 5(d) illustrates the change in IBV levels for the same groups under the 0.05um pore size membrane. Figure 5(e) highlights the change in Y17 Chlamydia strain levels for the same groups under the 0.002um pore size membrane; Figure 5(f) exhibits the change in IBV levels for the same groups under the 0.002um pore size membrane.

## 3 Discussion

Previous studies have demonstrated that Wolbachia is extensively distributed in arthropods and can induce host cytoplasmic incompatibility, parthenogenesis, and male killing. Recent research, however, has discovered that some Wolbachia strains can aid hosts in resisting viral infections, with certain strains exhibiting a broad-spectrum antiviral activity in their hosts. An example is the Wolbachia strain wAu in Drosophila melanogaster,, which exhibits high resistance against Drosophila C virus (DCV) and Flock House virus (FHV) in hosts^[26,29,53,54]^. Chlamydia, an obligate intracellular parasitic bacteria, is commonly found in vertebrates. As a microorganism situated between Rickettsia and viruses, Chlamydia shares a closer intracellular niche with viruses than Wolbachia does and is extensively present in vertebrates. We hypothesize that due to its fitness cost in hosts, it may assist the host in resisting related viruses, and earlier studies have demonstrated its potential antiviral effects^[14,21,41,46,55]^.

In this study, we aimed to isolate Chlamydia strains that can resist IBV using healthy chicken lung tissue samples from chicken farms, discovering that Chlamydia’s antiviral effect may be widespread. Our results indicate that 8 Chlamydia strains exhibit a significant inhibitory effect on IBV when B_i_ ≥ 0.800. Furthermore, cell co-cultivation observations revealed that it not only inhibited IBV but also AIV. This leads us to suspect that the isolated Y17 Chlamydia strain may exhibit a broad-spectrum antiviral effect in chickens. Our U-tube experiment showed that Chlamydia did not cause cells to secrete extracellular metabolites that could resist IBV. Instead, its antiviral effect was cell-autonomous, consistent with previous studies by Nainu et al. on Wolbachia^[52]^. Our U-tube experiments also demonstrated that its antiviral effect is unrelated to the host’s immune system. Notably, Slade et al.’s study showed that the antiviral effect of Cm wiess strain against HSV-2 does not work by stimulating host secretion of IFN-*β*^[15]^.

Our results suggest that isolating Chlamydia strains that antagonize IBV from healthy chickens on farms is a viable approach. The interaction between Chlamydia and IBV in the host may depend on the specific Chlamydia strain, and their relationship could be either synergistic or antagonistic^[14,21,31,32]^. These findings align with previous studies by Martinez et al. and Dodson et al. on the antiviral effect of Wolbachia in the host^[29,56]^. They showed that the interaction between related viruses and Wolbachia depends on the specific Wolbachia strain, but the antiviral effect is widespread in insects^[29,56]^. Our findings also help to explain the diverse relationships observed between Chlamydia and related viruses in previous studies, which could be synergistic, antagonistic, or unrelated, depending on the specific strain^[14,31]^. Overall, our results imply that isolating Chlamydia strains with antagonistic effects against IBV from healthy chickens in farms is feasible, and the antiviral effect of Chlamydia may be widespread, with some strains potentially possessing broad-spectrum antiviral effects.

Additionally, our results show that Chlamydia’s antagonistic effect on related viruses is cell-autonomous. Slade et al.’s study suggests that the antiviral effect of Cm is density-dependent, requiring viable Cm (or its components) to exist in the early stages of infection in the host without relying on stimulating the host to secrete antiviral IFN-*β* ^[14,15]^. Previous studies on the antiviral mechanism of Wolbachia in the host has demonstrated that its antiviral effect is also cell-autonomous and relies on the distribution density of Wolbachia in the host. This suggests that its antiviral effect is independent of the host immune response, and recent research indicates its antiviral effect is related to its early-stage impact on the virus genome structure (RNA). As a result, it can reduce the replication and infectivity of the infected virus in the host, appearing to change the genome structure of the offspring virus particles^[29,54,57,58]^. This antiviral mechanism is distinctly different from the previously discovered antagonistic effect of microorganisms, which primarily rely on the host’s immune system^[58,59]^. Consequently, it could represent a new and broad-spectrum antiviral mechanism for Wolbachia in insects. Based on the previous results of Slade et al. and our own study, the antiviral effect of Chlamydia also appears to be a new antiviral mechanism in the host^[14]^. We plan to further investigate these aspects in future study. Additionally, we aim to test whether the isolated Y17 Chlamydia strain has antagonistic effects on other respiratory viruses, such as Newcastle disease virus.

Previous studies have primarily focused on pathogenic Chlamydia strains, with most isolates being pathogenic. Our findings highlight the importance of studying non-pathogenic or low-toxicity Chlamydia strains^[38,39,40]^. We have shown that Chlamydia’s antagonistic effect against viruses may be widespread and that certain strains may have a broad-spectrum antiviral effect. Given that Chlamydia is widely present in vertebrates, including humans, and that most strains are non-pathogenic while only a few are pathogenic to the host, isolating these non-pathogenic Chlamydia strains and screening for antiviral strains, particularly broad-spectrum antiviral strains, may prove beneficial in preventing and treating vertebrate viral infections (including human infections)^[38,39,40]^. Furthermore, considering the previous studies conducted by Slade et al., Borel et al., and our own investigation, the antiviral effect of Chlamydia, similar to that of Wolbachia in insects, appears to constitute a new antiviral mechanism^[14,21]^. This warrants further investigation into the antiviral effect of Chlamydia, potentially leading to new ideas for the development of innovative antiviral methods or drugs. We plan to explore these areas in our future study.

## Supplementary

**Figure 1S.**
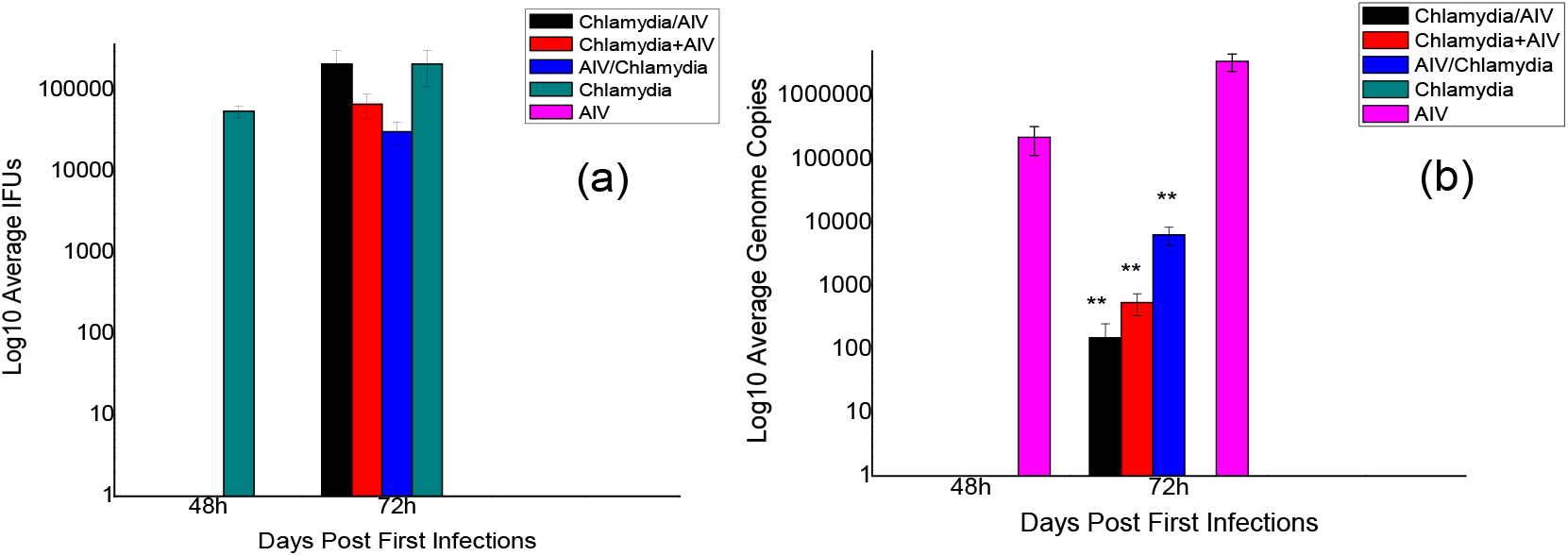
demonstrates the results of cell co-cultivation with Chlamydia and AIV in chicken tracheal epithelial cells. Figure 4(a) represents the changes in Chlamydia levels of groups, including Chlamydia/AIV, Chlamydia+AIV, AIV/Chlamydia, Chlamydia, and AIV. Figure 4(b) shows the changes in IBV levels of these same groups.

## Notes

### Competing Interest Statement

The authors have declared no competing interest.

### Summary of Updates

The section 1 and 2 has been revised.

