## Supplementary material for "Isolation of a Chlamydia Strain from Lung Tissue Samples in Healthy Chickens Exhibiting Antagonistic Effects on Avian Infectious Bronchitis Virus": Figure 1S

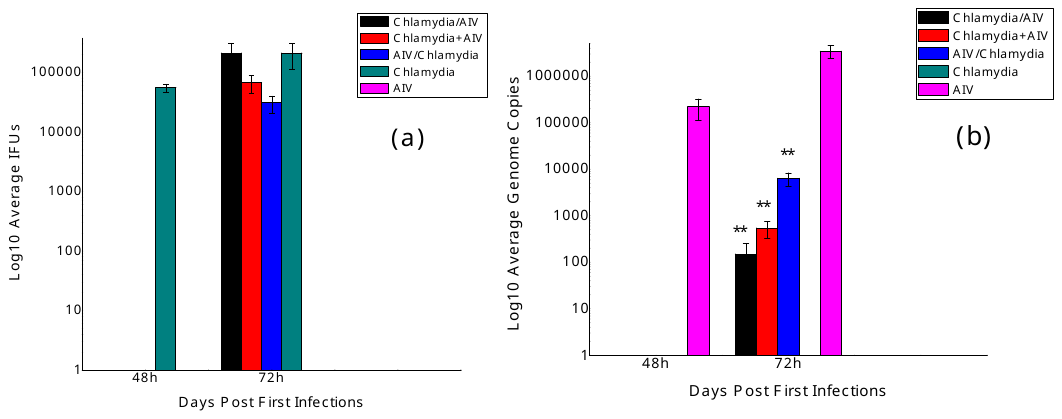

**Figure 1S** demonstrates the results of cell co-cultivation with Chlamydia and AIV in chicken tracheal epithelial cells. Figure 4(a) represents the changes in Chlamydia levels of groups, including Chlamydia/AIV, Chlamydia+AIV, AIV/Chlamydia, Chlamydia, and AIV. Figure 4(b) shows the changes in IBV levels of these same groups.
